# Aminoglycoside ribosome interactions reveal novel conformational states at ambient temperature

**DOI:** 10.1101/372144

**Authors:** Mary E. O’Sullivan, Frédéric Poitevin, Raymond G. Sierra, Cornelius Gati, E. Han Dao, Yashas Rao, Fulya Aksit, Halilibrahim Ciftci, Nicholas Corsepius, Robert Greenhouse, Brandon Hayes, Mark S. Hunter, Mengling Liang, Alex McGurk, Paul Mbgam, Trevor Obrinsky, Fátima Pardo-Avila, Matt Seaberg, Alan G. Cheng, Anthony J. Ricci, Hasan DeMirci

## Abstract

The bacterial 30S ribosomal subunit is a primary antibiotic target. Despite decades of discovery, the mechanisms by which antibiotic binding induces ribosomal dysfunction are not fully understood. Ambient temperature crystallographic techniques allow more biologically relevant investigation of how local antibiotic binding site interactions trigger global subunit rearrangements that perturb protein synthesis. Here, the structural effects of 2-deoxystreptamine (paromomycin and sisomicin), a novel sisomicin derivative, N1-methyl sulfonyl sisomicin (N1MS) and the non-deoxystreptamine (streptomycin) aminoglycosides on the ribosome at ambient and cryogenic temperatures were examined. Comparative studies led to three main observations. First, individual aminoglycoside-ribosome interactions in the decoding center were similar for cryogenic vs ambient temperature structures. Second, analysis of a highly conserved GGAA tetraloop of h45 revealed aminoglycoside-specific conformational changes, which are affected by temperature only for N1MS. We report the h44/h45 interface in varying states, that is, engaged, disengaged and in equilibrium. Thirdly, we observe aminoglycoside-induced effects on 30S domain closure, including a novel intermediary closure state, which is also sensitive to temperature. Analysis of three ambient and five cryogenic crystallography datasets reveal a correlation between h44/h45 engagement and domain closure. These observations illustrate the role of ambient temperature crystallography in identifying dynamic mechanisms of ribosomal dysfunction induced by local drug-binding site interactions. Together these data identify tertiary ribosomal structural changes induced by aminoglycoside binding that provides functional insight and targets for drug design.

## INTRODUCTION

The bacterial ribosome is a highly selected antibiotic target, with more than half of known antimicrobials targeting this enzyme (1,2). A hallmark example of ribosome targeting antibiotics is the aminoglycoside class of compounds, which were first discovered in the 1940s and deployed successfully in clinic before their ribosomal mechanism of action or toxicity to the human ear (ototoxicity) became clear (3-6). A prevailing model of drug-action suggests that drug interactions in the ribosomal A-site modulate antimicrobial potency and eukaryotic cellular dysfunction in the ear, hence further elucidation of the drug-ribosome structure activity relationship (SAR) is important for the design of safer antimicrobials with improved antibacterial activities (7).

Using the “diffract-before-destroy” strategy, serial femtosecond X-ray crystallography (SFX) enables the collection of high-resolution data at non-cryogenic temperatures using a new X-ray Free Electron Lasers (XFELs) (8). Study of the 4,6-deoxystreptamine aminoglycoside, paromomycin, at ambient temperature shows differences of up to 1.4 Å for the positions of several atoms at the drug-binding site in h44 between cryogenic and ambient temperatures(9). Here, we examine a 4,5-deoxystrepamine aminoglycoside, sisomicin, and novel less ototoxic N1-methyl-sulfonyl sisomicin derivative at cryogenic and ambient temperature **(Supplementary Fig. 1)** (10,11). For additional insight, we compare our data to previous paromomycin and streptomycin data sets, as well as the apo-30S state (12-15).

Aminoglycoside binding to the A-site in the 30S ribosomal subunit is a prominent theory of antibiotic action (13,16,17). In this model, aminoglycosides bind to the A-site in the 16S rRNA and cause the ribosome to misread pairings between tRNAs and mRNAs. A critical advance in our understanding of this interaction was made when high-resolution structures of 30S subunits complexed to aminoglycosides were attained, and when aminoglycoside-30S crystal structures were solved with the decoding complex (tRNA + mRNA) (14,15,18). In these structures, we can see that for 2-deoxystreptamine aminoglycosides ring I lies in the A-site helix and interacts with G1491 and A1493. This interaction forces adenines A1492 and A1493 into a flipped-out conformation which allows these adenines to form A-minor interactions with the first two Watson-Crick base pairs of the codon-anticodon mini-helix. This flipped out, so-called “locked” conformation forms the molecular basis for aminoglycoside induced mRNA misreading. Streptomycin, a non-deoxystreptamine aminoglycoside antibiotic, does not cause A1492 and A1493 flipping but acts as a wedge between the h44 and body of the 16S, thereby directly affecting the domain closure mechanism, also important in the miscoding process (see below)(15).

Local drug binding site interactions also trigger conformational changes further afield. In the apo-30S ribosomal subunit, a conserved 16S rRNA GGAA h45 tetra-loop exists in an equilibrium of engaged and disengaged states (15). Paromomycin binding to a 30S decoding complex structure shifts h45 towards the engaged state while streptomycin-binding shifts h45 towards the disengaged state (15). Additionally, during normal protein synthesis, two 30S subunit domains move closer to one another upon correct tRNA binding (14,17,19). Paromomycin binding stabilizes a closed domain conformation in 30S decoding complex, while streptomycin, locks the domain in an open state (15,17). To date, how aminoglycoside decoding center binding is connected to these events is unclear.

By probing co-crystals of sisomicin and N1MS at cryogenic and ambient temperatures, we aim to 1) determine if there are temperature-dependent differences in the local drug-binding site, 2) assess any detectable conformational changes in the 30S decoding complex, and 3) identify any common modes of action between these related chemical structures.

## MATERIALS AND METHODS

### Purification and crystallization of 30S ribosomal subunits for synchrotron cryo-crystallography

We purified and crystallized 30S ribosomal subunits from *T. thermophilus* HB8 (ATCC27634) (20) strain essentially as described (12,21). The mRNA fragment (with codon sequences underlined), 5’<u>UUU</u>UUU3’ and the anticodon stem loop ASL_Phe_, (with anticodon sequence underlined GGGGAUU<u>GAA</u>AAUCCCC, were purchased from Dharmacon (GE life sciences). 30S crystals (80μm × 80μm × 700μm) were sequentially transferred to the final buffer with 26% v/v 2-methyl-2,4-pentanediol (MPD) for cryoprotection and soaks were performed in the final buffer supplemented with 200μM of each mRNA oligos, ASL_Phe_ and 100μg/ml sisomicin parent compound or N1MS derivative compound for 48hours. Crystals were flash-frozen for data collection by plunging them directly into liquid nitrogen.

### Data collection and refinement of structure obtained by synchrotron X-ray cryo-crystallography

All synchrotron X-ray diffraction data were collected from a single crystal for all datasets. All datasets were collected with a Pilatus 6M detector at microfocus beamline BL12-2 at the Stanford Synchrotron Radiation Lightsource (SSRL) in Menlo Park, CA. Diffraction data sets were processed with the *XDS* package (22). Coordinates of the 30S subunit structure excluding mRNA and ASL (PDB accession code 4DR4) (15) with additional 30S rRNA and protein modifications were used for initial rigid body refinement with *PHENIX* (23) for all data sets. After simulated-annealing refinement, individual coordinates, three group B factors per residue, and TLS parameters were refined. Potential positions of magnesium or potassium ions were compared with those in a high-resolution (2.5 Å) 30S subunit structure (PDB accession code 2VQE) (24) in program *COOT* (25), and positions with strong difference density were retained. All magnesium atoms were replaced with magnesium hexahydrate. Water molecules located outside of significant electron density were manually removed. A similar refinement protocol was used for all datasets. The Ramachandran statistics for the parent compound (most favored / additionally allowed / generously allowed / disallowed) are 87.7 / 11.6 / 0.7 / 0.0 % respectively. The Ramachandran statistics for the parent compound (most favored / additionally allowed / generously allowed / disallowed) are 87.4 / 11.9 / 0.8 / 0.0 % respectively. The structure refinement statistics are summarized in **Supplementary Table 1**. Structure alignments were performed using the alignment algorithm of *PyMOL* (www.schrodinger.com/pymol) with the default 2σ rejection criterion and five iterative alignment cycles. All X-ray crystal structure figures were produced with *PyMOL* <u>_ENREF_9</u>.

### Preparation and Crystallization of 30S Ribosomal Subunits SFX crystallography at XFEL

30S ribosomal subunits from *T. thermophilus* HB8 (ATCC27634) (20) were prepared as previously described (12,21). Purified 30S ribosomal subunits were crystallized at 4°C by the hanging drop method using a mother liquor solution containing 17% (v/v) 2-methyl-2,4-pentanediol (MPD), 15 mM magnesium acetate, 200 mM potassium acetate, 75 mM ammonium acetate and 100 mM 2−(*N*-morpholino) ethanesulfonic acid (MES)-KOH (pH 6.5). Microcrystals 2−3 × 2−3 × 3−4 μm^3^ in size were harvested in the same mother liquor composition, pooled (total volume of 500 μl loaded) and supplemented with 200μM of each mRNA oligos, ASL_Phe_ and 100μg/ml sisomicin parent compound or N1MS derivative compound 48 hours before data collection. Crystal concentration was approximated to be 10^10^−10^11^ particles per ml based on light microscopy and nanoparticle tracking analysis (NanoSight LM10-HS with corresponding Nanoparticle Tracking Analysis (NTA) software suite (Malvern Instruments, Malvern, UK).

### coMESH construction

The central sample line carrying 30S ribosomal subunit microcrystals was a continuous 100 μm inner diameter fused silica capillary with a 160 μm outer diameter and a length of 1.5 m (Polymicro, Phoenix, AZ, USA). This capillary directly connected the reservoir with the interaction region, passing through a vacuum feedthrough and then through a microfluidic *Interconnect Cross* (C360-204, LabSmith, Livermore, CA USA; C360-203 *Interconnect Tee* may also be used with no need to plug the fourth channel) (**Supplementary Figure 2**). Feeding the sample through an uninterrupted straight capillary without filters, unions or connections minimized the risk of sample clogs and leaks, which most often occur at flow channel transitions. In addition, the large capillary inner diameter allowed the crystal slurry to flow in the system with no inline filtration or filters at the reservoir. The fitting at the top of the *cross* in **Supplementary Figure 2** was an adapter fitting (T116-A360 as well as T116-100, LabSmith, Livermore, CA, USA) which compressed a 1/16” (1.6 mm) OD, 0.007” (178 μm) ID polymer (FEP) tubing sleeve (F-238, Upchurch, Oak Harbor, WA, USA) onto the sample capillary. This connection method was necessary to properly seal the capillary to the *cross*. The sheath line at the bottom of the figure was a short 5 cm fused silica capillary with an outer diameter of 360 μm and an inner diameter of 180, 200, or 250 μm depending on the desired flow rate of the sheath liquid possible with the driving pressure available. The length and ID of this line were chosen to allow sufficient flow for the sheath line. This outer capillary was connected to the *cross*, and the sheath liquid was supplied from the third port of the cross (the left port in the figure) through a compatible microfluidic tubing; we typically used silica capillaries with 360 μm outer diameters and 100 μm inner diameters for optimum flow rate. For simplicity, the injector has been designed to operate at relatively low backing pressures (up to a few atmospheres). For this experiment, the sheath liquid line was connected to a syringe filled with the appropriate sister liquor, driven by a syringe pump (PHD Ultra, 703006, Harvard Apparatus, Holliston, MA, USA) We electrically charged the sheath liquid between 0-5,000 V potential using an in-line conductive charging union (M-572, Upchurch, Oak Harbor, WA, USA) connected to a SRS PS350 (Stanford Research Systems, Sunnyvale, CA, USA) high voltage source.

The capillary assembly was loaded into the CXI 1 × 1 μm^2^ focused beam chamber(26) with a custom load lock system developed at Lawrence Berkeley National Laboratory. A grounded, conical counter electrode with a 1 cm opening was placed approximately 5 mm below the capillary tip; the capillaries and opening of the cone were coaxial. The angle of the cone and its distance from the tip were set to enable a diffraction cone with a 45° half-angle. All capillaries were fed through vacuum flanges with 1/16” (1.6 mm) Swagelok bulkhead fittings using appropriately sized polymer sleeves. The sheath reservoir was a 1 ml Gastight Hamilton syringe with a PTFE Luer tip (1001 TLL SYR, Hamilton, USA). The ribosome microcrystalline sample was supplied from a 500 μl Hamilton gas tight syringe.

### Operating the coMESH

Before connecting the central sample line, the sister liquor was loaded, flowed and electrically focused. Once a stable jet was achieved, the central sample line carrying ribosome slurry was connected. Notably, the sister liquor never fully stabilized because of the entrained air from the disconnected sample line continuously introducing bubbles. The central sample line had much less fluidic resistance compared to the outer line; connecting it first with a vacuum sensitive sample will cause immediate jet clogging and blockages, for this reason the outer line should always be on while operating in vacuum environment. The sister liquor was set to flow at, or near, the flow rate of the mother liquor. If diffraction hits were not being observed, the sister liquor flow was reduced and/or the mother liquor flow was increased slowly to ensure the jetting remains stable, and until the diffraction patterns appear.

### Selecting a sister liquor for ribosome microcrystalline slurry

The main goal of the sister liquor is to survive the vacuous environment (10^−5^ Torr) and protect the inner sample line from the adverse vacuum effects. In the case of the co-terminal coMESH (9), the compatibility between the sister liquor and the ribosome crystals was not considered since the fluid interaction occurred just before the sample is probed, allowing minimal time, if any, for the fluids to mix. The limiting constraint is typically the availability of the fluid and its viscosity. For simplicity, the 26% MPD-containing sister liquor seems to be an ideal starting choice. The buffer has proven to be quite reliable and survives the vacuum injection with minimal issues of leaving behind dried precipitate such as salt or PEG to stay adhered to the capillary tip. The lower viscosity allows the sister liquor to be pumped easily by a standard syringe pump through the more resistive *cross* manifold or concentric annular flow before the exit region of the capillary.

### Sample Injection

#### coMESH for ribosome crystals

The sample reservoir was loaded with 30S ribosome crystal slurry in mother liquor, described above. The sister liquor was identical to the mother liquor in that the original substituent concentrations remained constant with the exception of having a higher MPD concentration of 26% (v/v). The sample capillary was a 100 μm ID × 160 μm OD × 1.5 m long fused silica capillary. The sheath flow capillary was a 75 μm ID × 360 μm OD × 1 m long fused silica capillary. The outer concentric capillary was a 200 μm ID × 360 μm OD × 5 cm long tapered silica capillary. The tips of the inner and outer capillaries were located at the same position. The taper of the outer capillary minimizes the surface on which debris can build up on after X-ray interaction with the sample but is not necessary for operation. The applied voltage on the sheath liquid was typically 3,000 V, and the counter electrode was grounded. The sample ran typically between 0.75-1 μl/min and the sheath flow rate typically matched the sample flow or was slightly faster.

For the study of 30S-sisomicin and 30S-N1MS complex crystals, the inner sample line contained unfiltered crystals in their native mother liquor containing 17% (v/v) 2-methyl-2,4-pentanediol (MPD). The size distribution of the 30S crystals was uniform (2-3 × 2-3 × 4-5 μm^3^) due to their controlled slower growth at 4°C (27). Occasional larger sized 30S crystals were discarded by repeated gentle differential settling without centrifugation. The outer sister liquor was the same buffer, with no crystals, as the mother liquor in that the original substituent concentrations remained constant while having increased the MPD to 26% (v/v) to aid in vacuum injection. The coMESH injector allowed us to successfully deliver 30S ribosomal subunit complex crystals (**Supplementary Figure 2**) and collect sufficient data to solve a complete ambient-temperature SFX structures, which was previously not attainable at an XFEL (27).

### Data Collection and Analysis for SFX studies at LCLS

#### Diffraction data collection of 30S ribosomal subunit microcrystals

The microcrystals were screened at SSRL BL12-2 in preparation of LCLS beamtime. The SFX experiments with 30S ribosome microcrystals were carried out at the LCLS beamtime ID: cxil1416 (N1MS derivative) and cxim7916 (sisomicin parent compound) at the SLAC National Accelerator Laboratory (Menlo Park, CA). The LCLS X-ray beam with a pulse duration of 50 fs was focused using X-ray optics in a Kirkpatrick-Baez geometry to a beam size of 1.3 × 1.3 μm^2^ full width at half maximum (FWHM) at a pulse energy of 2.9 mJ, a photon energy of 9.5 keV (1.29 Å) and a repetition rate of 120 Hz.

A total of 1,533,919 and 1,336.966 detector frames were collected with the Cornell-SLAC Pixel Array Detector (CSPAD)(28) from sisomicin and N1MS respectively. The total beamtime needed for sisomicin and N1MS dataset were 213 and 185 minutes respectively, which shows the efficiency of this injector system, as due to lack of blockages no dead time was accumulated. Individual diffraction pattern hits were defined as frames containing more than 30 Bragg peaks with a minimum signal-to-noise ratio larger than 4.5, which were a total of 290,894 and 285,745 images for sisomicin and N1MS respectively. The detector distance was set to 223 mm, with an achievable resolution of 3.08 Å at the edge of the detector (2.6 Å in the corner).

After the detection of hits and following the conversion of individual diffraction patterns to the HDF5 format, the software suite *CrystFEL* (29) was used for crystallographic analysis. The peak location information from *CHEETAH* (30) was used for the indexing of individual, randomly oriented crystal diffraction patterns using FFT-based indexing approaches. The detector geometry was refined using an automated algorithm to match found and predicted peaks to subpixel accuracy (31). The integration step was performed using a built-in Monte-Carlo algorithm to estimate accurate structure factors from thousands of individually measured Bragg peaks (32). After the application of per pattern resolution cutoff, frames which didn’t match to an initial merged dataset with a Pearson correlation coefficient of less than 0.4, were excluded from the final dataset. For the sisomicin the final set of indexed patterns, containing 25,401 frames (8.7% indexing rate), was merged into a final dataset (Overall CC* = 0.97; 3.4 Å cutoff) for further analysis (P4_1_2_1_2, unit cell: a = b = 402.6 Å, c = 175.9 Å; α = β = γ = 90°). The final resolution cutoff was estimated to be 3.4 Å using a combination of *CC* ^34^* and other refinement parameters. The final dataset had overall *R_split_* = 33.3%, and *CC\** = 0.52 in the highest resolution shell. For the N1MS the final set of indexed patterns, containing 23,956 frames (8.4% indexing rate), was merged into a final dataset (Overall CC* = 0.96; 3.5 Å cutoff) for further analysis (P4_1_2_1_2, unit cell: a = b = 402.1 Å, c = 175.1 Å; α = β = γ = 90°). The final resolution cutoff was estimated to be 3.5 Å using a combination of *CC* (33*) and other refinement parameters. The final dataset had overall *R_split_* = 51.0%, and *CC\** = 0.45 in the highest resolution shell.

### Ambient temperature 30S Ribosomal Subunit SFX Structure refinement

We determined the ambient temperature 30S□sisomicin parent compound and 30□N1MS derivative complex structures using the automated molecular replacement program *PHASER* (34) with the previously published cryo-cooled 30S ribosomal subunit□mRNA□ASL□paromomycin complex synchrotron structure as a search model (PDB entry 4DR4) (15). The resulting structure was refined with rRNA modifications, which are mostly located near the decoding center at which sisomicin and N1MS binds. This choice of starting search model minimized experimental variations between the two structures such as sample preparation, crystal growth, model registry and data processing. Coordinates of the 4DR4 with additional RNA and protein modifications were used for initial rigid body refinement with the *PHENIX* (23) software package. During initial refinement of ambient temperature XFEL structure, the entire mRNA and ASL components were omitted and the new models were built into unbiased difference density. After simulated-annealing refinement, individual coordinates, three group B factors per residue, and TLS parameters were refined. Potential positions of magnesium or potassium ions were compared with those in a high-resolution (2.5 Å) 30S subunit structure (PDB accession code 2VQE)(24) in program *COOT* (35), and positions with strong difference density were retained. All magnesium atoms were replaced with magnesium hexahydrate. Water molecules located outside of significant electron density were manually removed. The Ramachandran statistics for parent compound (most favored / additionally allowed / generously allowed / disallowed) are 88.5 / 10.8 / 0.7 / 0.0 % respectively. Ramachandran statistics for N1MS derivative (most favored / additionally allowed / generously allowed / disallowed) are 88.9 / 10.5 / 0.6 / 0.0 % respectively. The structure refinement statistics are summarized in **Supplementary Table 1**. Structure alignments were performed using the alignment algorithm of *PyMOL* (www.schrodinger.com/pymol) with the default 2σ rejection criterion and five iterative alignment cycles. All X-ray crystal structure figures were produced with *PyMOL.*

### Conformational analysis of the structural ensemble

The ensemble studied is listed in **Supplementary Table 3**.

### Pre-processing

Each structure was pre-processed using PyMOL. When two alternate chains were built, each was extracted separately and two resulting models were built (resulting in the engaged/disengaged states for example). Chains were renamed according to the following nomenclature. Chain A: 16S rRNA, residues numbered 1 to 1532. Chains B to U were kept intact: they constitute the 20 proteins bound to the 16S rRNA. Chain V: tRNA ASL in the P site. The symexp command was used to generate the closest symmetry mate within 20 Å from nucleotide 1493 in chain A, and the anticodon stem loop mimicked by the 16S rRNA protruding in the P-site of the decoding center was extracted and built in the model. The remaining was discarded. Chain W: tRNA ASL in the A site, when present. Chain X: ligand molecule akin aminoglycoside antibiotic, when present. Chain Y: mRNA moiety, when present. Chain Z and w, ions and water molecules. The analysis was carried out on chain A only, further reducing it to one atom per nucleotide (Phosphorus). When a given nucleotide is absent from at least one structure, it is trimmed away from all others, thus resulting in a consistent set of 12 structures made of the same 1504 atoms. The RMSD between all pairs in that set was computed and is given in **Supplementary Table 2**.

### Principal Component Analysis

The 12×1504 matrix *A* containing the structural set was decomposed (SVD) as a product of 3 matrices *USV^T^*: *U* is a 12×1504 matrix containing the left singular vectors of *A*, *S* is a 12×12 diagonal matrix containing the singular values of *A* and *V* is a 12×12 matrix containing the right singular vectors of *A*. The vectors in *U* are also called the principal component vectors of the ensemble, sorted in decreasing order of their associated singular value. The latter are related to the percentage of the ensemble variance explained by the given component: the larger the singular value, the more the associated component explains the diversity in the ensemble. *V* informs on how components are mixed for each individual in the set; it is used to compute the projection of individual structures along a given component (see **Supplementary Table 3**).

### Visualization

The first component vector field is depicted with arrows in **Fig. 3a** using the module *modevectors* in PyMOL, and as a periodic movie in **Supplementary Movie 1** with home-made code that interpolates a periodic function between the ensemble average structure and its first component vector.

## RESULTS

### Cryogenic temperature sisomicin-30S decoding complex

To study the structural impact of sisomicin binding to the bacterial ribosome, we crystallized the small 30S ribosomal subunits of *Thermus thermophilus* as previously described (12,21). We then prepared the decoding complex containing sisomicin, hexauridine mRNA, and a phenylalanine ASL (ASL_Phe_) by soaking these compounds into 30S crystals (**see Methods**). A complete X-ray diffraction data set was collected at beamline 12-2 of Stanford synchrotron radiation lightsource (SSRL) from a single, large cryo-cooled crystal after 24-hour soaking. We determined the crystal structure at 3.4 Å resolution by merging four complete 360-degrees rotation datasets collected at 5% X-ray transmission from different regions of the same rod-shaped crystal (**Supplementary Table 1**). A clear positive difference density in the unbiased electron density map was visible in the binding site associated with sisomicin (**Fig. 1a**). Sisomicin binds to h44 of the 30S-decoding center by forming seven hydrogen bonds and a hydrophobic ring stacking **(Supplementary Fig. 3)**. Specifically, ring-I establishes base stacking with decoding center residue G1491, and ring-II engages in H-bonding interactions with backbone phosphate of 16S rRNA residue A1493 and bases of G1494 and U1495. Ring-III is involved in a single H-bonding with backbone phosphate of U1406, two H-bonds with base G1405 and single H-bond with the 5-methyl modified cytosine (5mC) base 5mC1407 **(Supplementary Fig. 3)** (*Escherichia coli* numbering is used throughout).

**Figure 1.**
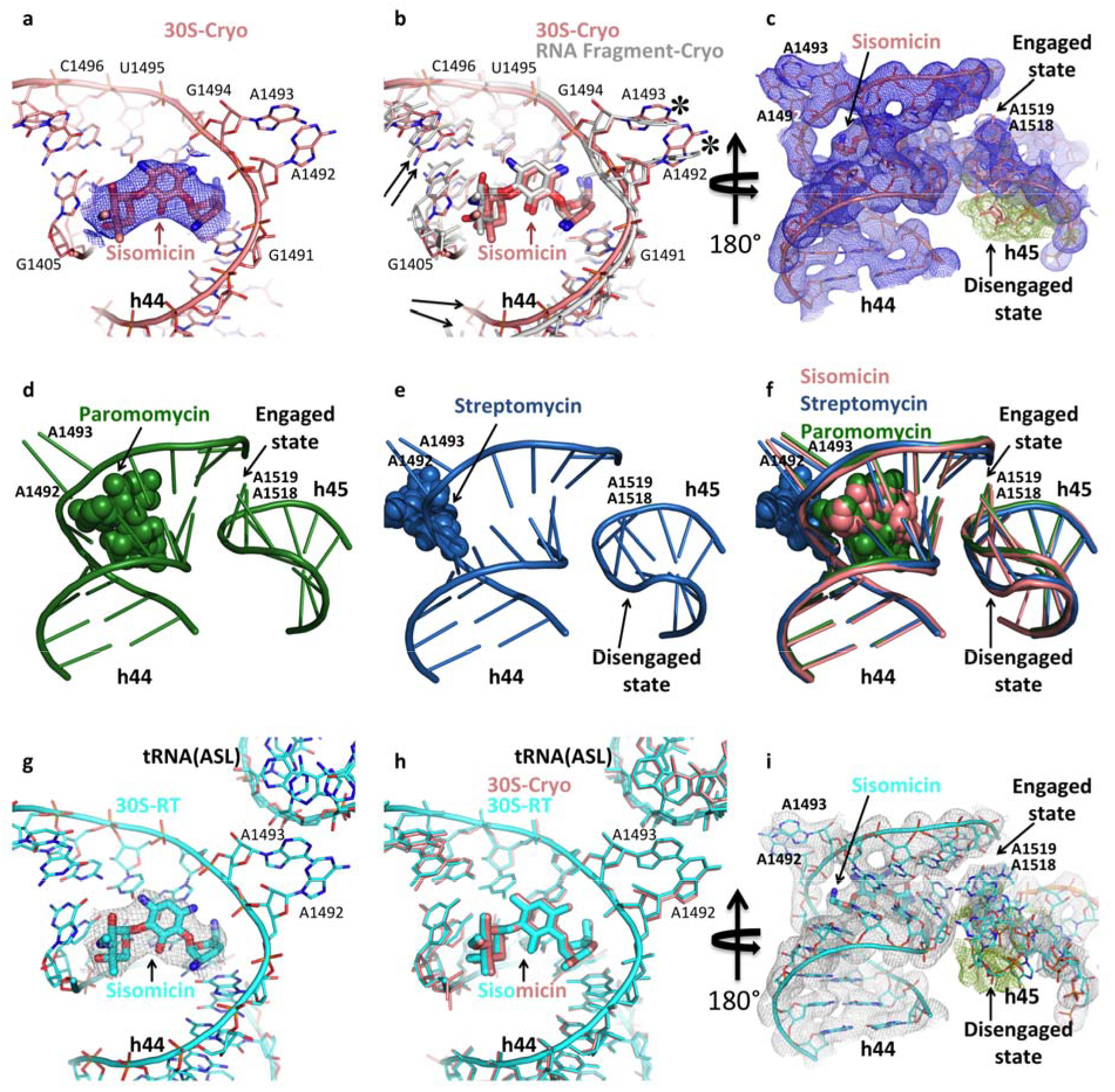
Aminoglycosides-decoding center structures at cryogenic (first and second rows) and ambient temperature (third row). (**A)** Sisomicin in the decoding center in a cryogenic sisomicin-30S decoding complex structure (PDB ID: 6CAP). Unbiased Fo-Fc simple difference electron density map that belongs to sisomicin shown in blue mesh and contoured at 3σ level. **(B)** Superposition of the decoding center from a cryogenic sisomicin-30S decoding complex structure (salmon) and a cryogenic short RNA fragment-bound sisomicin structure (gray) (PDB ID: 4F8U). Black arrows show minor local conformational changes in the 16S rRNA backbone. Asterisks mark positions of A1492 and A1493. (**C)** The h44-45 interaction in the cryogenic sisomicin-30S decoding complex is in an engaged: disengaged equilibrium. 2Fo-Fc electron density map of the h44-45 helices contoured at 1.5σ level and colored in blue. The unbiased Fo-Fc simple difference electron density map of the h45 region contoured at 3σ level and colored in green, which indicates the presence second alternate conformation. **(D)** The h44-45 interaction in the cryogenic paromomcyin-30S decoding complex is fully engaged (PDB ID: 4DR4). **(E)** The h44-45 interaction in the cryogenic streptomycin-30S decoding complex is fully disengaged (PDB ID: 4DR6). **(F)** Superposed sisomicin-, paromomycin-and streptomycin-30S decoding complex structures. Sisomicin stabilizes a novel aminoglycoside-induced conformation by maintaining h45 in an equilibrium state (salmon). **(G)** Sisomicin in the decoding center of ambient temperature sisomicin-30S decoding complex structure (PDB ID: 6CAR). Unbiased Fo-Fc electron density of the sisomicin calculated from ambient temperature SFX diffraction data shown in gray mesh and contoured at 3σ level. The structure will be colored in cyan throughout. **(H)** Superposition of the ambient (cyan) and cryogenic (salmon) sisomicin-30S decoding complex structures. **(I)** The h44-45 interaction in the ambient temperature sisomicin-30S decoding complex structure is in an engaged: disengaged equilibrium. 2Fo-Fc electron density map of the h44-45 helices contoured at 1.5σ level and colored in gray. The unbiased Fo-Fc electron density map of the h45 region contoured at 3σ level and colored in green indicates the presence of second alternate conformation.

In the absence of mRNA and ASL, sisomicin-30S complex crystals diffracted poorly to below 4.5 Å resolution. Despite the low resolution of sisomicin-30S structures, A1492 and A1493 were observed flipped out from h44 (data not shown). In the absence of mRNA and ASL, the addition of streptomycin (a non-flipping aminoglycoside) (15) improved the resolution of sisomicin-30S structures to 3.75 Å. In these structures, A1492 and A1493 were flipped out into open conformation suggesting that sisomicin binding was sufficient to stabilize the flipped conformation. Independent of streptomycin, we also determined that the addition of an mRNA and ASL improved the resolution of sisomicin-30S structures to 3.4 Å. Hence, sisomicin-30S and N1MS-30S X-ray diffraction data was collected from crystals with a decoding complex. The crystal structure of the sisomicin-30S decoding complex adopted the near-canonical decoding conformation, with the h44 residues A1492 and A1493 flipped out of h44, consistent with other aminoglycosides (17).

To compare the impact of sisomicin binding on the bacterial ribosome, we superposed our 30S decoding structure with the only sisomicin structure available in the Protein Data Bank (4F8U): sisomicin in complex with a short RNA oligonucleotide dimer (36) (**Fig. 1b**). The observed root-mean-square deviation (RMSD) between 16S rRNA atoms of our structure and the published oligomer-drug complex of 1.23 Å suggests the local minor conformational differences exist in the backbone of the 16S rRNA between our data and the published structure. The positions of A1492 and A1493, which engage in direct hydrogen-bonding interaction with the first and second position of the codon-anticodon helix, were very similar overall (indicated by asterisks) Similar deviations up to 2 Å of the positions of A1492 and A1493 backbone atoms in the different cryogenic 70S ribosome structures with cognate A-site tRNAs were also observed (37,38) (**Fig. 1b**).

Previous studies demonstrate that aminoglycosides alter the interaction between residues on h44 and a conserved tetraloop (G1516-A1519) on h45 (15,39). The h44-45 interaction involves the dynamic formation and breaking of a network of H-bonds between h44 (U1406, G1496 and G1497 residues) and the neighboring h45 (G1517, A1518 and A1519 residues) resulting in engaged and disengaged conformations of h44-45(15,21,40,41). In the apo-30S structure h45 exists in an engaged: disengaged equilibrium (15). Here, sisomicin binding favors the equilibrium state of the h44-45 interaction in the decoding complex (**Fig. 1c**). This observed aminoglycoside-stabilized state is novel in that paromomycin binding favors the engaged conformation, and streptomycin binding favors the disengaged conformation (**Fig. 1d-f**)(15).

### Ambient temperature sisomicin-30S decoding complex

To evaluate potential differences in the crystal structures at ambient temperature, co-crystals of the sisomicin-30S decoding complex were probed in an SFX experiment at the Coherent X-ray Imaging (CXI) instrument of the Linac Coherent Light Source (LCLS; Menlo Park, CA, USA) by using a low-flow concentric electrokinetic liquid injector setup **(Supplementary Figure 2)** (9,26). During the protein screening beamtime, a 3.4 Å resolution complete dataset was collected from 2−3 × 2−3 × 3−4 μm^3^ size microcrystals supplemented with sisomicin, mRNA and ASL_Phe_ oligonucleotide complex. Total of 1,533,919 images collected with 290,894 hits and 25,401 indexed patterns during 213 minutes of net data collection time. The resulting electron density map indicated similar drug density quality in the unbiased difference electron density maps (**Fig. 1g**). The crystal structure of the sisomicin-30S decoding complex at ambient temperature adopted the near-canonical decoding conformation, with the h44 residues A1492 and A1493 flipped out toward the minor groove of ASL and mRNA pair, consistent with the cryogenic data (**Fig. 1h**).

Analysis of dynamic interactions at the h44-45 interface revealed that at ambient temperature sisomicin binding favors an equilibrium state of h44-45 engagement: disengagement **(Fig. 1i)**. This is consistent with the apo-30S structure and the cryogenic sisomicin-30S decoding complex data (**Fig. 1c, i**). The minimal binding differences observed between the cryogenic and ambient temperature sisomicin-30S structures (**Fig. 1c,i**) suggest that ribosomal decoding complexes bound to this antibiotic can be probed at cryogenic temperature and may still be representative of what occurs at ambient temperature.

### Cryogenic temperature N1MS-30S decoding complex

Structural insights into the sisomicin-30S decoding complex provide a basis against which novel derivative compounds can be compared. We examined co-crystals of the N1MS-30S decoding complex following the procedures described above. The N1MS-30S decoding complex structure was determined to 3.4 Å resolution. Comparative analysis of the N1MS-and sisomicin-30S decoding complex structures obtained at cryogenic temperatures revealed the drug binding with the decoding center to be markedly similar, with an overall RMSD of 0.34 Å (**Fig. 2a,b**). The overall decoding state, including A1492 and A1493 extra-helical flipping and local hydrogen-bonding network of the drug-30S subunit, was maintained. Investigation of dynamic interactions at the h44-45 interface revealed the interaction to be in an engaged: disengaged equilibrium (**Fig. 2c**). This is identical to the one that is observed in the sisomicin-30S decoding complex crystal structure (**Fig. 1c**).

**Figure 2.**
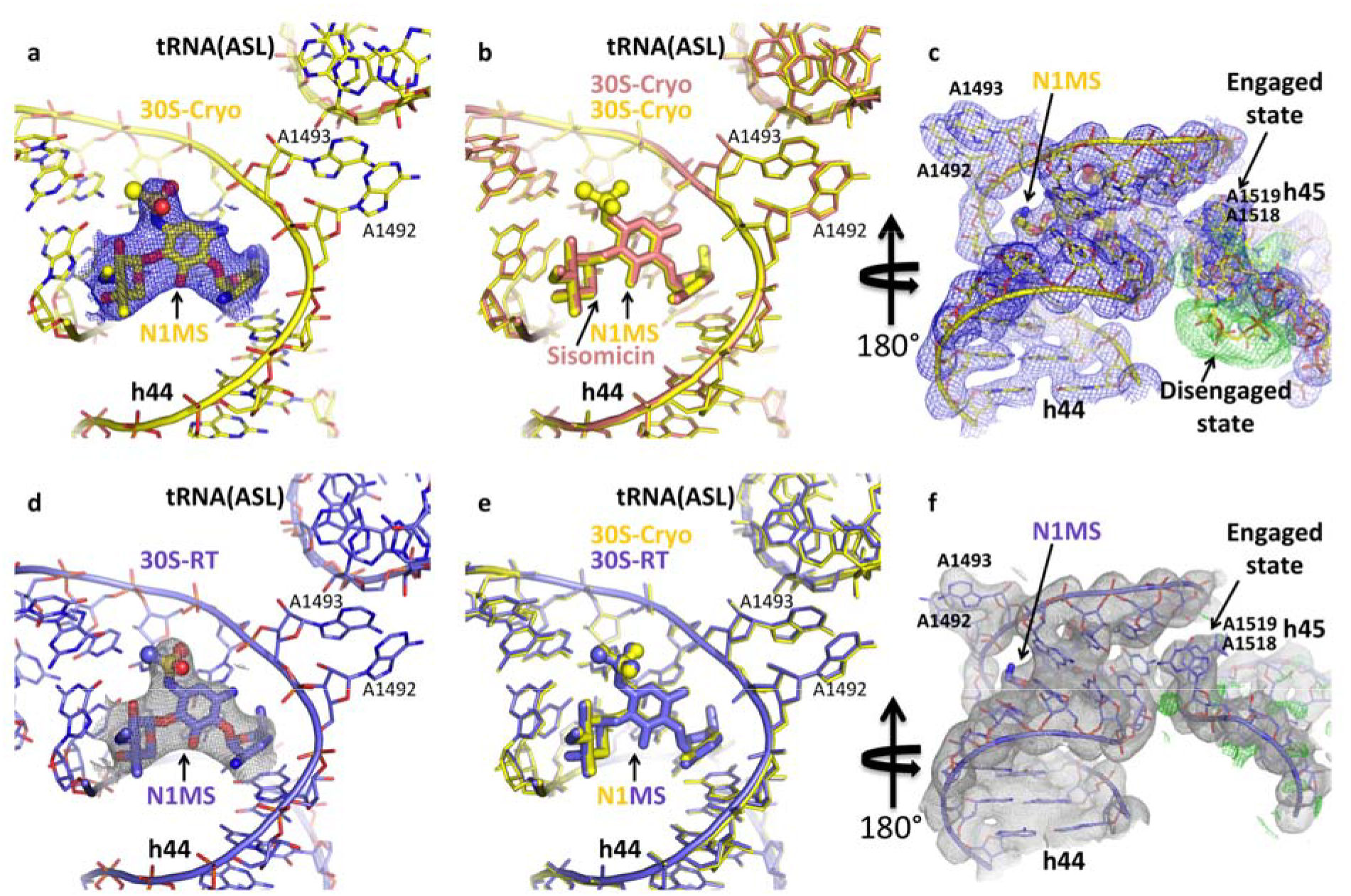
N1MS in the decoding center at cryogenic (first row) and ambient temperature (second row). **(A)** N1MS in the decoding center of cryogenic N1MS-30S decoding complex structure (PDB ID: 6CAQ). Unbiased Fo-Fc electron density of the N1MS calculated from cryogenic temperature diffraction data shown in blue mesh and contoured at 3σ level. The structure will be colored in yellow throughout. **(B)** Superposition of the cryogenic sisomicin-30S decoding complex structure (salmon) and the cryogenic N1MS-30S decoding complex structure complex (yellow). **(C)** The h44-45 interaction in the cryogenic N1MS-30S decoding complex is in an engaged: disengaged equilibrium. 2Fo-Fc electron density map of the h44-45 helices contoured at 1.5σ level and colored in blue. The unbiased Fo-Fc electron density map of the h45 region contoured at 3σ level and colored in green indicates the presence of second alternate conformation. **(D)** N1MS in the decoding center of ambient temperature N1MS-30S decoding complex structure (PDB ID: 6CAS). Unbiased Fo-Fc electron density of the N1MS derivative compound calculated from ambient temperature diffraction data shown in gray mesh and contoured at 3σ level. The structure will be colored in slate throughout. **(E)** Superposition of th ambient (slate) and cryogenic (yellow) N1MS-30S decoding complex structure. **(F)** The h44-45 interaction in the ambient N1MS-30S decoding complex is in a fully engaged state. 2Fo-Fc electron density map of the h44-h45 helices contoured at 1.5σ level and colored in gray. The unbiased Fo-Fc electron density map of the h45 region contoured at 3σ level and colored in green indicates the absence of second alternate conformation that is observed at cryogenic temperature shown in (c).

### Ambient temperature N1MS-30S decoding complex

To evaluate potential differences between ambient temperature N1MS-30S and sisomicin-30S structures, co-crystals of the same ribosomal decoding complex bound to N1MS were then probed in an SFX experiment at the CXI instrument **(Supplementary Figure 2)**. During a second protein screening beamtime, a 3.5 Å resolution complete dataset was collected from 2-3 × 2-3 × 3-4 μm^3^ size microcrystals supplemented with N1MS, mRNA and ASL_Phe_ oligonucleotides complex (**Fig. 2d**). Total of 1,336,966 images collected with 285,745 hits and 23,956 indexed patterns during 185 minutes of net data collection time.

At ambient temperature, the binding interactions of N1MS in the decoding state were similar to the cryogenic structure with a decoding site RMSD of 0.40 Å (**Fig. 2d,e**). The resulting electron density maps were high quality at 3.5 Å resolution, and the positive difference density that belongs to methylsulfonyl modification was clearly observed in the simple unbiased difference omit density map (**Fig. 2d**). Surprisingly, analysis of the h44-45 interface in the N1MS-30S decoding complex structure revealed an alternate conformation at ambient temperature. The h45 tetraloop appeared locked in a fully engaged conformation at ambient temperature while at cryogenic temperature h45 was in an engaged: disengaged equilibrium. Thus, the h44-45 interaction in the N1MS-30S structure shows clear differences at ambient and cryogenic temperatures and represents the first temperature-dependent dynamic conformational changes reported in this region (**Fig. 2c,f**).

### Sisomicin and N1MS induces novel domain closure states

To gain further structural insights, we performed global scale analysis of the 30S subunit by superposing the apo-30S subunit structure with sisomicin-and N1MS-30S decoding complex structures obtained at cryogenic and ambient temperatures, together with structures previously described, either bound to streptomycin decoding complexes at cryogenic temperature, or bound to paromomycin decoding complexes at both cryogenic and ambient temperature (15,42). This allowed us to better understand the impact of sisomicin and N1MS on the large scale motions of the ribosome, in particular, domain closure in the 30S subunit – an important conformational change that coincides with recognition of the correct codon-anticodon interaction (14). The degree of domain closure is considered binary, being either in open or closed states during normal ribosome decoding (14). In previous aminoglycoside-related studies, paromomycin was shown to stabilize domain closure, while streptomycin was shown to favor a more open arrangement of the shoulder relative to the rest of the 30S (15,17). We decomposed our structural ensemble into principal components (**see Methods**). Most of the structural variability was explained with a component whose main feature is to bring the shoulder domain toward the platform. The head domain follows an anti-correlated motion, resulting in a contraction at the decoding center (**Fig. 3a,b** and **Supplementary Movie 2**), akin to the domain closure motion previously described (17). Projection of all individual structures in our ensemble onto this principal component allows us to order them according to the extent of open/closed states of the shoulder domain (**Fig. 3c-e** and **Supplementary Table 3**). The resulting ordering is consistent with the apo-30S state being the most open, the streptomycin-bound state being almost as open as the apo-30S state, while paromomycin-bound structures being the most closed. Interestingly, the sisomicin-and N1MS-30S decoding complex structures suggest that a novel intermediate domain-closure state exists (**Fig. 3e**). Both sisomicin and N1MS datasets indicate that these drugs stabilize the domain closure in a less-closed state somewhere between paromomycin and streptomycin. Since both sisomicin and N1MS are antimicrobial our structural data suggest that the partial domain closure states observed do not hinder aminoglycoside-induced ribosomal dysfunction, but may be part of their antimicrobial mechanism (11).

**Figure 3.**
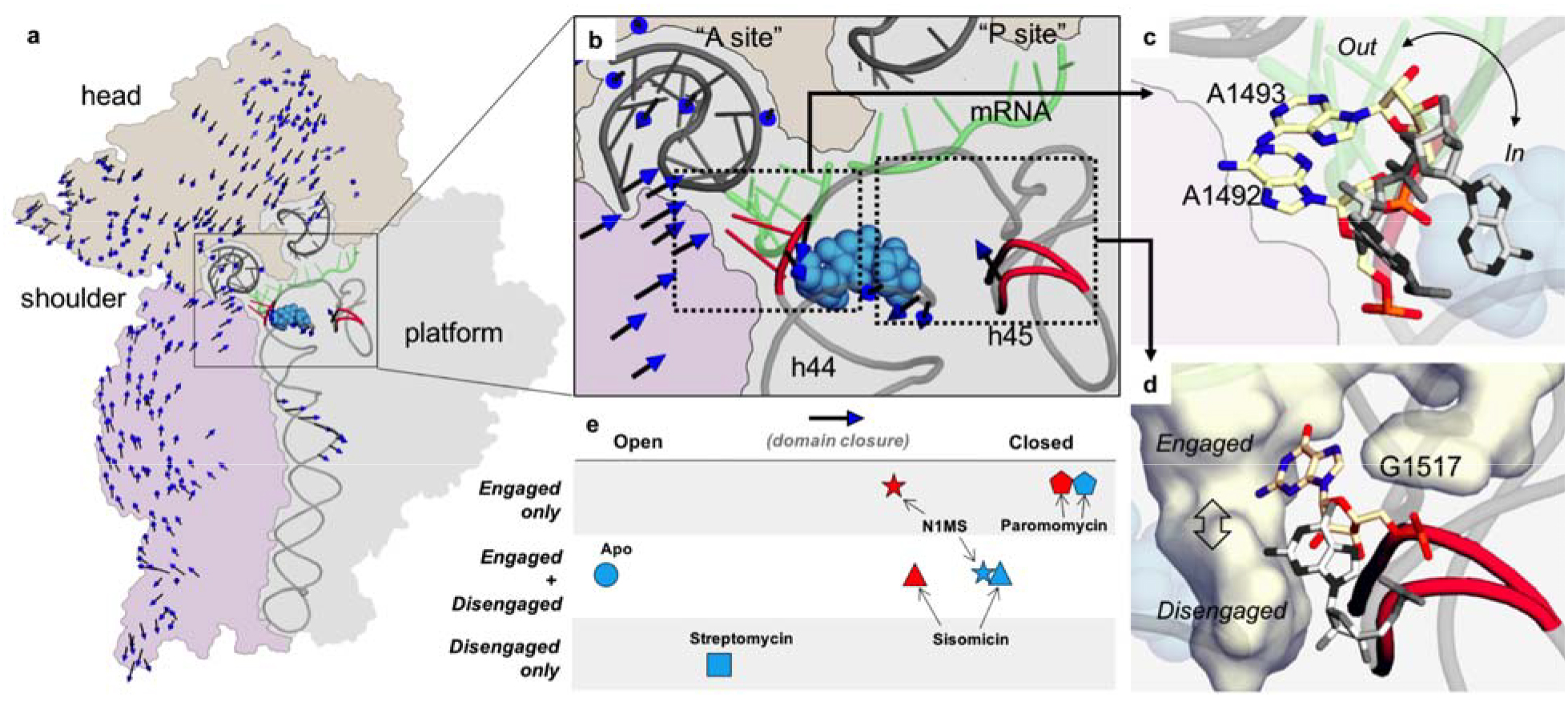
Sisomicin and N1MS binding to the 30S decoding complex induces an intermediate state of 30S domain closure. **(A)** The 30S ribosomal subunit illustrating the location of the drug binding pocket and the site of the h44-45 interactions. The drug is shown with blue spheres. The 30S subunit contour is shown with three different colored regions of interest, and the portion of the rRNA making helices h44 and h45, the ASL_Phe_ and mRNA moieties are shown as gray ribbons. The first principal component (**see Text**) is displayed as a vector field from the ensemble average structure. Note that smaller displacements were filtered out, and the arrows displayed are arbitrarily scaled, for clarity. More local motions of interest are further highlighted in red in the decoding center. **(B)** Detailed view of the decoding center displaying the drug binding pocket, where A1492 and A1493 (red sticks in left box) are displaced in response to aminoglycoside binding. The two h45 states, engaged and disengaged are also shown on the right (red ribbon in right box). **(C)** Detailed view of the flipping bases, in the context of the decoding center where the A-site tRNA (top left, gray), the mRNA (green) can be seen close to the outward flipped bases, and the aminoglycoside molecule (blue sphere) is seen superposed with the inward flipped state of the same bases. **(D)** Engaged and disengaged conformations of h45. The pocket in which G1517 is engaged, made in part by h44, is shown as a surface. Proximity to the drug (spheres in the background) binding site is apparent. **(E)** Ordering of the apo-30S (circle) structure and the streptomycin-(square), sisomicin-(triangle), N1MS-(star) and paromomycin-(pentagon) 30S decoding complex structures. Blue represents cryogenic structures, red represents ambient temperature structures studied along the 1st principal component extracted from their conformational ensemble. The vertical scale shows h45 disengagement, engagement or equilibrium in the corresponding crystal structure.

## DISCUSSION

Bacterial protein synthesis is a major antibiotic target in the cell, and structural investigations into the aminoglycoside-ribosomal SAR provide a valuable framework on which to develop new compounds.

### Role of cryogenic data in drug design

Recent paromomycin-30S structures showed differences in the drug-binding site between cryogenic and ambient temperatures(9). Unlike paromomycin, the local drug-binding pattern is similar for both parent and designer sisomicin compounds at both temperatures (Fig. 1h, 2b, and 2e). These observations collectively suggest there are instances in which local binding pocket interactions of crystal structures determined under cryogenic conditions reflect conformations also observed at ambient temperature, thus indicating cryo-crystallography can serve as a reliable higher-throughput antibiotic-ribosome experimental screening platform to investigate binding. The ability to ascertain these details routinely in a relevant model system such as 30S subunit *in vitro* can help accelerate the design and identification of viable aminoglycoside derivative candidates. Current scarcity, cost and availability of XFEL beamtime preclude the routine use of ambient temperature SFX crystallography.

### Novel h44/45 states at cryogenic and ambient temperatures

Our investigation of sisomicin and N1MS at both cryogenic and ambient temperature reveals novel states. At cryogenic temperatures, sisomicin and N1MS do not perturb the h45 engaged: disengaged equilibrium that exists in the apo-30S state (Figs. 1c, 2c)(15). This is surprising since previous studies report aminoglycosides to alter this equilibrium state (**Fig. 3e**)(15,41). Moreover, we observe this equilibrium state shifts to engaged for the N1MS compound with temperature but not for sisomicin.

Why N1MS, but not sisomicin, has a temperature-dependent effect on the h44/h45 interaction is likely related to the N1-methyl sulfonyl modification. On analysis of the N1MS A-site binding, we propose a new hydrogen bond may be forming between the N1-methyl-sulfonyl modification and C1496, similar to the bond that forms between the N1-Haba modification on amikacin and C1496 (43,44). This new interaction may alter C1496 hydrogen bonding with methyl groups on h44 A1518 and A1519 in the GGAA tetraloop, these hydrogen bonds are part of a network that stabilize the packing interactions between h44 and h45. Future higher resolution structures and molecular dynamics simulations may reveal a relationship between the N1 methyl-sulfonyl modification and the 16S rRNA backbone that alters the energy thresholds associated with the structural dynamics of h45 disengagement, engagement and equilibrium.

The location of h44/h45 activity points to a critical role in bacterial ribosome function. The sequence of the GGAA h45 tetraloop is highly conserved, as is the m^6^_2_A methylation modification on A1518 and A1519, and the m^6^_2_A modification enzyme itself, the KsgA methyltransferase (39,45-47). In bacteria, deletion of KsgA and lack of m^6^_2_A methylation, results in a translational dysfunction, a cold-sensitive phenotype and resistance to kasugamycin. Kasugamycin is a non-deoxystreptamine aminoglycoside with a binding site different than the 2-deoxystreptamine subset studied here (48-50). We hypothesize that its resistance mechanism operates through the rescue of the h44/h45 dynamics impaired by the lack of methylation of h45, thus possibly sharing a modulation role of the h44/h45 activity with the aminoglycosides studied in this work. This site also appears to have a fundamental role in eukaryotic systems. Knockout of the mouse mitochondrial equivalent of the KsgA methyltransferase (mitochondrial transcription factor B1 - TFB1M) and lack of methylation is embryonically lethal, and genetic variants in the human *TFB1M* has been associated with diabetes and aminoglycoside related hearing loss (51-54).

### Novel domain closure state

Here, we report a previously unreported “intermediary” state of domain closure, a conformation that challenges the binary open and closed mode currently proposed in the literature (Ogle et al., 2002). Domain closure (shoulder movement and head rotation) normally coincides with tRNA selection and is linked to activation of Elongation Factor Tu (EF-Tu), an enzyme that hydrolyses GTP and releases the amino-acyl tRNA in the A-site so translation can proceed (17). We hypothesize that the intermediary state may be due to the chemical differences between the drugs studied and propose that this novel conformation which may be a signature of the 4,6-disubstituted 2-deoxystreptamine subset of aminoglycosides.

To identify relationships between local A-site binding and non-local conformational changes we investigated correlations between A-site binding, h44-45 engagement and domain closure, and report a correlation between h45 engagement and domain closure (**Fig. 3e**). While further studies are required, this correlation could potentially explain kinetic data indicating that aminoglycosides have different ribosomal mechanisms of action (e.g. streptomycin and paromomycin)(55). Streptomycin is reported to lock the ribosome in a structure that is unable to respond normally to the binding of a cognate tRNA thus lowering the rate and/or fidelity of translation, whereas paromomycin is suggested to switch the ribosome into a high affinity and high-activity conformation that accelerates GTP hydrolysis, thus lowering the accuracy of tRNA selection (55).

In conclusion, we describe previously unreported states of h45 and domain closure specific to the 4,6-disubstituted 2-deoxystrepatmine aminoglycoside antibiotics. In particular, we report a temperature sensitive effect on the 30S ribosomal subunit induced by N1MS. To the best of our knowledge this is also the first-time A-site binding has been linked to global 30S subunit rearrangements through h44/45 interactions – a result that not only underscores the importance of ambient temperature crystallography but also demonstrates how small modifications to A-site binding drugs can induce dramatically different ribosomal subunit conformations.

## ACCESSION NUMBERS

Protein Data Bank: Coordinates and structure factors of cryogenic Sisomicin-30S, cryogenic 30S-N1MS, ambient temperature Sisomicin-30S and ambient temperature N1MS-30S complexes have been deposited under accession codes 6CAP, 6CAQ, 6CAR and 6CAS respectively.

## SUPPLEMENTARY DATA

Supplementary Data are available at NAR Online.

## ACKNOWLEDGMENT

The authors would like to thank Dr. Michael Levitt for his support and critical reading of the manuscript. Portions of this research were carried out at the Linac Coherent Light Source (LCLS) and Stanford Synchrotron Radiation Lightsource (SSRL) at the SLAC National Accelerator Laboratory. LCLS and SSRL are supported by the U.S. Department of Energy (DOE), Office of Science, Office of Basic Energy Sciences (OBES) under Contract No. DE-AC02-76SF00515. The SSRL Structural Molecular Biology Program is supported by the DOE Office of Biological and Environmental Research, and by the National Institutes of Health, National Institute of General Medical Sciences (including P41GM103393). Parts of the sample injector used at LCLS for this research was also funded by the National Institutes of Health, P41GM103393, formerly P41RR001209. The LCLS is acknowledged for beam time access under experiment no. cxil1416 (N1MS derivative), cxim7916 (sisomicin parent compound). EHD, RGS and HD acknowledge the support of the OBES through the AMOS program within the CSGB and the DOE through the SLAC Laboratory Directed Research and Development Program. EHD acknowledges financial support from the Stanford University Dean of Research. HD acknowledges support from the joint Stanford ChEM-H and SLAC National Accelerator Laboratory seed grant program and also NSF Science and Technology Centers grant NSF-1231306 (Biology with X-ray Lasers, BioXFEL). CG acknowledges SLAC and the DOE for financial support through the Panofsky fellowship. This work is supported by the National Institute of Health (NIH) grant R01DC014720 (AJR and AGC). FP, NC and FPA acknowledge National Institute of Health (NIH) grant R35GM122543 to (Michael Levitt). We thank A. Fry’s continuous support for providing LCLS intern contributions to the portions of the experiments carried at LCLS.

## AUTHOR CONTRIBUTIONS

HD and AJR designed and coordinated the project. HD, EHD, YR, HC, FA prepared the samples for SFX and synchrotron studies. RGS, HD, AM, PM, BH, TO and EHD build the coMESH injectors. FP, CG executed the SFX data reduction. NC and FPA provided realtime XFEL data analysis. HD refined ribosome structures. The experiments were executed by HD, RGS, FP, NC, FPA, MH, ML, MS, FA, HIC, YR, EHD, and MOS. Data were analyzed by HD, FP MOS, AJR, and AGC. The manuscript was prepared by HD, EHD, FP, MOS, and AJR with input from all the coauthors.

